# Selection drives the evolution of convergent gene expression changes during transitions to co-sexuality in haploid sexual systems

**DOI:** 10.1101/2021.07.05.451104

**Authors:** Guillaume G. Cossard, Olivier Godfroy, Zofia Nehr, Corinne Cruaud, J. Mark Cock, Agnieszka Lipinska, Susana M. Coelho

## Abstract

Co-sexuality has evolved repeatedly from ancestors with separate sexes across a wide range of taxa. The switch to co-sexuality is expected to involve major molecular readjustments at the level of gene expression patterns, as modified males or females will express the opposite sexual function for which their phenotypes have been optimized. However, the molecular changes underpinning this important transition remain unknown, particularly in organisms with haploid sexual systems such as bryophytes, red and brown algae. Here, we explore four independent events of emergence of co-sexuality from uni-sexual (dioicous) ancestors in brown algal clades in order to examine the nature, evolution and degree of convergence of gene expression changes that accompany the breakdown of dioicy. The amount of male versus female phenotypic differences in dioicous species were not correlated with the extent of sex-biased gene expression, in strike contrast to what is observed in animals. Although sex-biased genes exhibited a high turnover rate during brown alga diversification, their predicted functions were remarkably conserved. Transition to co-sexuality consistently involved adaptive gene expression shifts and rapid sequence evolution, particularly of male-biased genes. The gene expression profiles of co-sexual species were more similar to those of females than to males of related dioicous species, suggesting that the former may have arisen from ancestral females. Finally, we identified extensive convergent gene expression changes associated with the transition to co-sexuality, and these changes appear to be driven by selection. Together, our observations provide novel insights on how co-sexual systems arise from ancestral, haploid UV sexual systems.

## Introduction

Eukaryotic organisms exhibit a wide diversity of sexual systems, ranging from separate sexes (referred to as gonochorism in animals and dioecy in plants) to co-sexuality (combined sexes), and several theories have been developed to explain what conditions favour which strategy (Ghiselin, 1969; Charnov et al., 1976; Maynard Smith, 1978; Charnov, 1982; Charlesworth, 1999, 2006; Barrett, 2002; Vamosi et al., 2003; Jarne & Auld, 2006; Meagher, 2007). The evolution of this diversity often involved transitions between sexual systems. For example, separate sexes have evolved from co-sexual ancestors independently many times in several eukaryotic lineages, and the fundamental mechanisms and evolutionary drivers of this important transition have been intensively studied in many organisms (reviewed in ^1,2^). Frequently, organisms with separate sexes display marked sexual dimorphism in a range of morphological, behavioral and physiological traits. Females and males are nevertheless genetically similar with the exception of the sex-specific regions of their sex chromosomes. While sex-chromosomes necessarily play a role in the expression differences between sexes, most of sex-biased gene expression involves autosomal genes ^3–5^. Differences in autosomal gene expression patterns between sexes may be associated with different physiological processes directly linked to the production of male or female gametes (primary sexual dimorphism) or to the consequences of sexual selection and/or sexual specialization (secondary sexual dimorphism) that may occur once separate sexes have evolved ^6^.

While the emergence of separate sexes from co-sexual ancestors and the evolution of sexual dimorphism have been thoroughly investigated ^5,7–9^, less attention has been devoted to the opposite transition, i.e. from separate sexes to co-sexuality. Transitions to co-sexuality have occurred frequently during eukaryotic evolution and are relatively common in animals (e.g. Denver et al, 2011; Avise & Mank, 2009; reviewed in Weeks, 2012) and land plants (e.g. Lloyd, 1975; Pannell, 1997 Schaefer and Renner, 2010; reviewed in Renner, 2011). In flowering plants this transition was believed to be rare but recent studies are increasingly providing evidence that dioecy-to-monoecy transitions may have occurred frequently ^10,11^. Evolutionary models intending to decipher the causes of such transitions invoke the sex-allocation theory ^12^ and the deterministic fate of genetic modifiers causing the acquisition of an opposite-sex function ^13,14^. However, empirical knowledge on the proximate mechanisms and forces driving the shift from separate sexes to co-sexuality remains largely elusive.

Transitions from separate sexes to co-sexuality are also prevalent in eukaryotic lineages other than animals and flowering plants, and in particular those that express sex during the haploid stage of their life cycles. In organisms such as bryophytes, liverworts, green, red and brown algae, male and female sexes are expressed during the haploid (gametophyte) stage ^15^. Genetic sex determination in organisms with haploid sexual systems occurs during meiosis (and not at fertilisation as in XY and ZW systems) ^16^, depending on whether spores inherit a U or V chromosome ^17,18^. Spores receiving a V sex chromosome will develop into a male individual (male gametophyte) and the spores inheriting a U sex chromosome will grow into females (female gametophytes). Organisms with haploid sex determination may also display epigenetic sex determination (so called monoicy), where an haploid, co-sexual, individual gametophyte may produce both male and female sexual structures ^19,20^. Despite the prevalence of haploid sexual systems among eukaryotes, the mechanisms underlying transitions from dioicy to monoicy are so far unknown.

In this context, the brown algae represent a particularly attractive group for studies of the evolution of sexual systems and breakdown of dioicy. The brown algae are part of the stramenopile (or heterokont) supergroup, which also includes diatoms and oomycetes, and they have diverged from the Archaeplastida lineage at the time of the eukaryotic crown radiation ^21^. Most brown algae have a haplo-diplontic life cycle, with a haploid gametophyte generation alternating with a diploid sporophyte generation. In these brown algae, sexuality is expressed in the haploid generation, with male and female gametes either produced by the same (monoicy) or on two separate individuals (dioicy). Dioicy is the prevalent reproductive system ^19,22^. This situation contrasts markedly with that described for flowering plants, where only about 6% of extant species have separate sexes and is more similar to that of bryophytes and liverworts ^20^. Dioicous brown algae may exhibit a broad range of levels of sexual dimorphism, both at the level of the gametophytes but also between male versus female gametes size ^19,22^. While the predicted ancestral state in the brown algae is dioicy, transitions to monoicy have occurred frequently and independently in the different clades ^22,23^. The independent emergence of monoicous lineages from dioicous ancestors makes this group particularly interesting to examine the genomic consequences and mechanisms underlying the breakdown of dioicy.

Here, we explore multiple, repeated events of loss of dioicy (Figure 1) to investigate the molecular basis and level of convergence of the shifts to co-sexuality. We demonstrate a lack of correlation between phenotypic sexual dimorphism and gene expression levels among dioicous brown algae. Ancestral state reconstruction indicated high turnover rates of sex-biased genes, yet independently recruited sex-biased genes shared similar functions across the species. We then focused on changes in gene expression patterns of orthologous genes that are specifically or preferentially expressed in haploid males and females, when they function in a monoicous context. Male-biased genes were particularly concerned by both adaptive expression shifts and faster evolutionary rates associated with the transition to monoicy. Monoicous species displayed expression profiles that were more similar to those of the female of the closely related dioicous species than to the male. Finally, we identified a striking amount of convergent gene expression changes associated with the emergence of co-sexuality, which were likely driven by selection.

**Figure 1.**
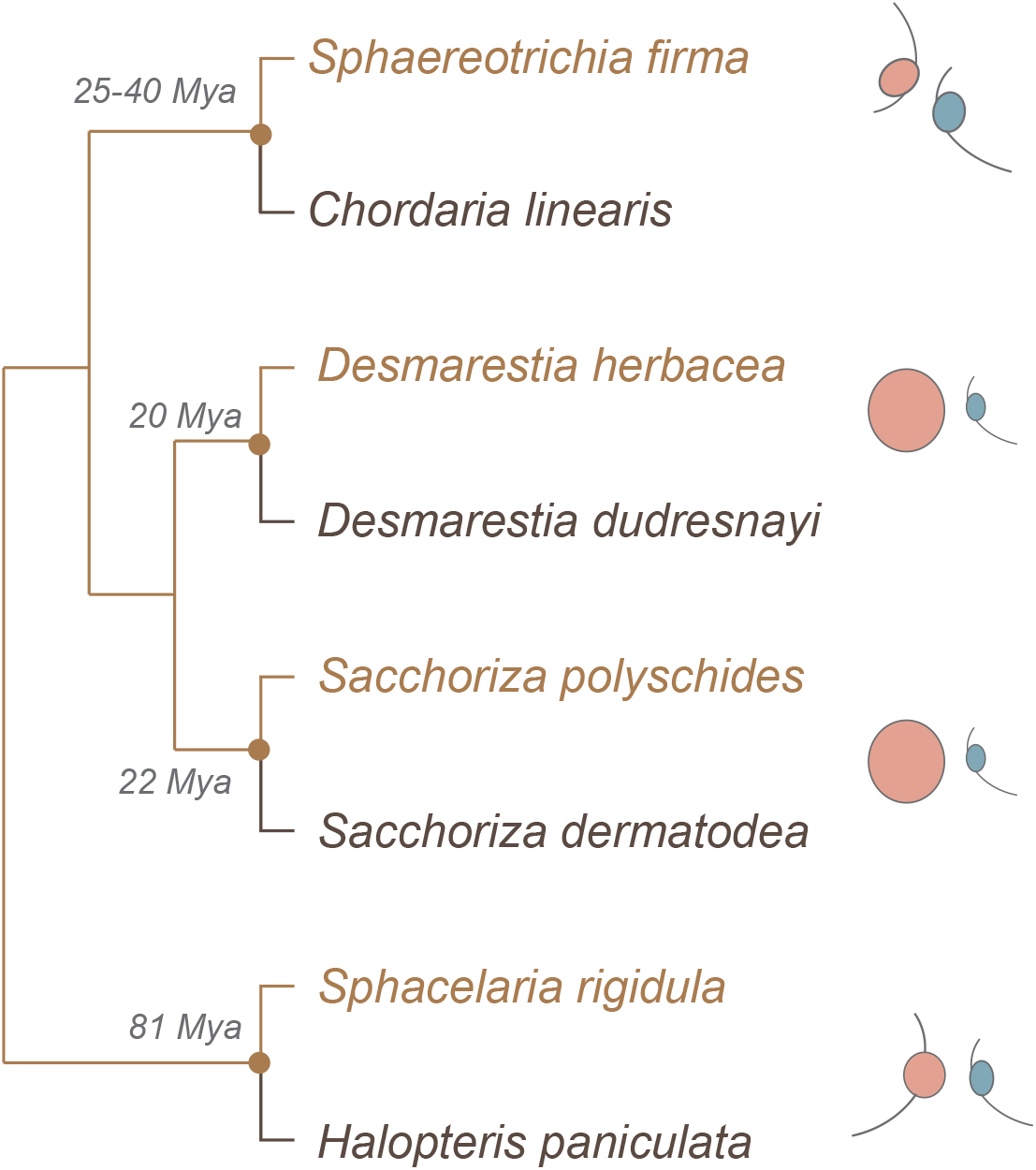
Diagram of the phylogeny of the eight species of brown algae investigated. Approximate estimated age of nodes is based on (Kawai et al., 2015); O. de Clerck pers. communication). A schematic view of typical gamete size differences (female in red, male in blue) per species pair is presented.

## Results

The present study examines sex-biased gene expression in dioicous brown algae and the gene expression changes associated with the transition from dioicy to monoicy. We based our analysis on transcriptomes sequenced from pairs of dioicous-monoicous species in four major clades of brown algae spanning app. 200 million years of evolution ^24^. The transitions are predicted to have occurred at different times in the past (between 20 and 88 MY; Figure 1). Each pair represents an independent transition from dioicy to monoicy. We chose dioicous species with different levels of gamete dimorphism, reflecting the diverse levels of gamete dimorphism occurring across brown algae.

### Sex-biased gene expression in dioicous brown algae

Gene expression patterns in gametophytes of the eight brown algal species were measured by deep sequencing (RNA-seq) of cDNA from male, female and co-sexual gametophytes. Transcript abundance (measured as transcripts per million, TPM) was strongly correlated between biological replicates with *r*^2^ ranging from 0.89 to 0.99 (Table S1). Counts of expressed genes (TPM>5^th^ percentile counts across all genes in at least two samples) identified a number of expressed genes that ranged from 13,180 to 27,391 (Figure S2, Table S1).

Deseq2 was used to identify genes that were differentially expressed in each of the sexes of the dioicous species ^25^. The analysis retained only genes that displayed at least a 2-fold change expression level between sexes (FC>2, p_adj_<0.05). Note that sex-linked genes (genes located in the sex-specific regions on the V (male) and U (female) sex chromosomes; see methods), were removed from the set of sex-biased genes and thus excluded from further analysis.

All four dioicous brown algae displayed substantial sex-biased gene expression, at least compared with plants and other brown alga ^7,9,26^ ranging from 12.7 % of the expressed genes in *S. rigidula* to 33.3% in *S. firma* (Figure 2A-2B, Table S2). We found similar proportions of male-biased compared with female-biased genes for the majority of the studied species (Figure 2A-2B) with the exception of *S. polyschides*, where male-biased genes were more abundant than female-biased genes (15.81% male-biased genes versus 9.82% female-biased genes; Chi^2^-test *P* < 2.2 x10^−16^).

**Figure 2.**
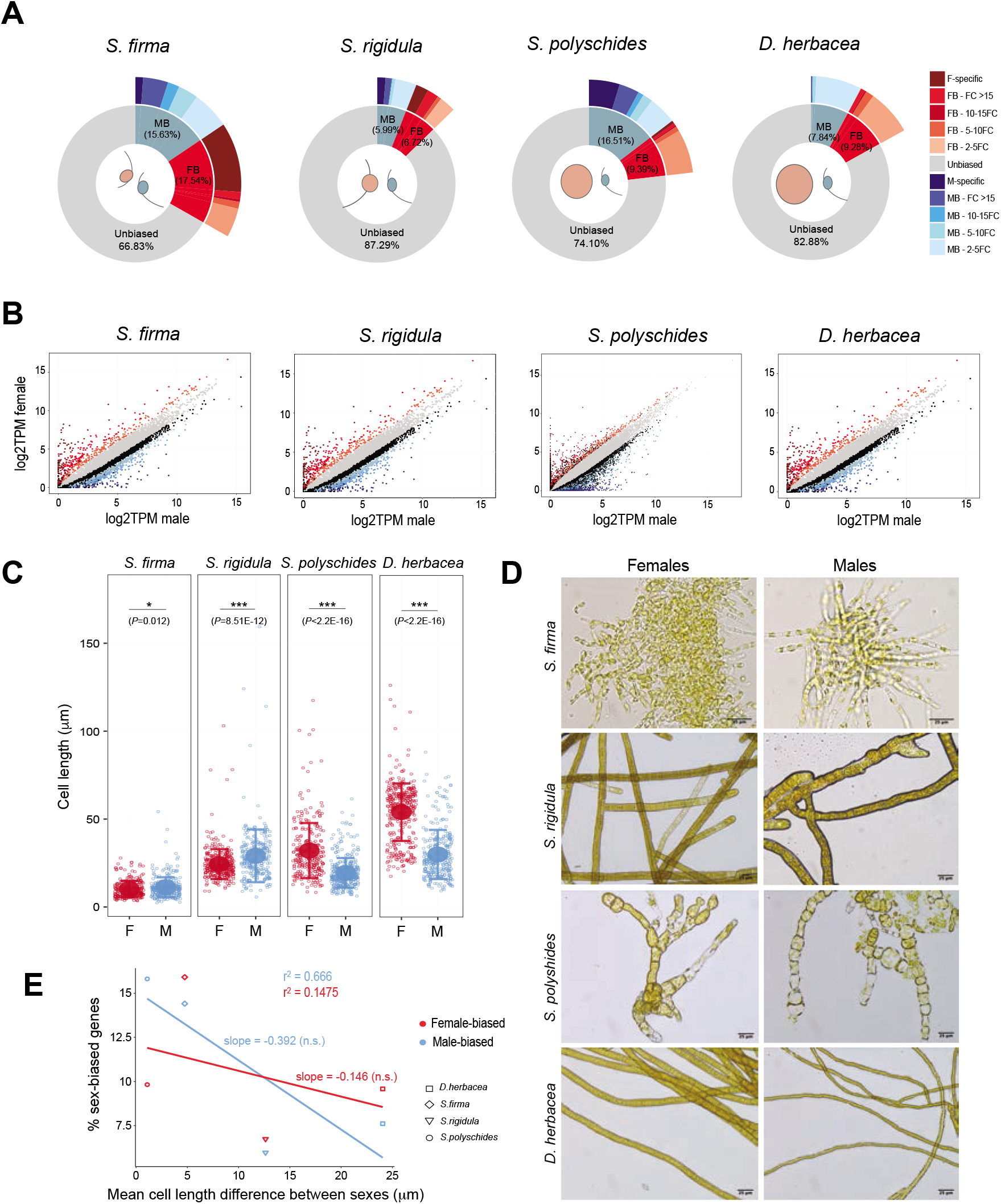
Patterns of sexual dimorphism in dioicous brown algae. A) Pie charts representing the fractions of sex-biased genes among expressed genes (female-bias in red, male-bias in blue) in the four dioicous species. Gradients of colors represent the intensity of expression fold-change (FC), from 2FC difference to more than 15FC. The percentages are calculated based on the total number of expressed genes averaged across sexes. B) Comparison of gene expression levels, in log2(TPM+1), between males and females within dioicous species. Colour patterns follow the ones used in panel A, except for black points which represent unbiased genes that presented a FC > 2. C) Scatterplots of the lengths of cells of immature gametophytes of dioicous species. The mean and standard deviations are plotted per sex per species. Stars indicate significant difference between mean cell length, tested with t-tests. *0.01 < P < 0.05; **0.001 < P < 0.01; ***P < 0.001. D) Micrographs of male and female immature gametophytes viewed under an inverted light microscope for each dioicous species investigated. E) Linear regressions of the fraction of female- and male-biased genes (in red and blue, respectively) among expressed genes against the mean difference in cell length recorded between the sexes (in μm), in the four dioicous species investigated.

### Sex-biased gene expression and phenotypic sexual dimorphism

To investigate the link between sex-biased gene expression and the level of sexual dimorphism, we carried out morphometric measurements of male and female gametophytes complemented with literature searches. These measurements allowed us to quantify the amount of phenotypic dimorphism present in each of the four dioicous species (Table S3, Figure 2C). In all dioicous species, gamete size dimorphism was coherent with sexual differences in terms of gametophyte cell size (Table S3). For example, *D. herbacea* gametophytes presented marked sexual dimorphism both at level of gamete size and gametophyte cell length, whereas *S. firma* was the species with least sexual difference both in terms of gametophyte morphology and gamete size (Table S3, Figure 2C-D).

In animals, sexual differences at the phenotypic level are correlated with levels of sex-biased gene expression ^8,27^, but this correlation has not been found in plants ^26^. We compared the differences in gametophyte cell size between males and females with the proportion of sex-biased genes in each of the four dioicous brown algal species. We detected no correlation between phenotypic sexual dimorphism (gametophyte cell size) and the number of sex-biased genes (Figure 2E). For instance, *S. firma* was the species that exhibited the highest level of sex-biased gene expression and nonetheless presented the lowest level of phenotypic sexual dimorphism.

Taken together, our observations indicate a considerable level of sex-biased gene expression in the four dioicous species studied here, but the level of sex-biased gene expression did not reflect the level of morphological dimorphism between males and females.

### Evolution of sex-biased gene expression in dioicous species

We next investigated how sex-biased gene expression has evolved by comparing the four dioicous brown algal species. Orthofinder identified a total of 14,017 orthogroups (OGs), of which 2,098 contained only one gene per species and therefore represented the set of 1:1:1:1 OGs. An additional 2,778 OGs had a single member in each of three of the studied species (i.e., the gene was missing in the fourth species). We considered that these 1:1:1:0 OGs, which likely represent single copy ancestral genes that were lost in one of the species, also provide useful information about conservation of sex-biased gene expression. Note that the 1:1:1:0 OGs could also represent OGs where one of the genes is missing from one of the genome assemblies, particularly the draft genome assembly (*S. rigidula*). Furthermore, we also included 1,085 orthogroups with a duplicated gene in a single species that aligned along more than 60% of their length (Figure 3A), resulting in 5,961 ‘dioicous single-copy orthologs’ (DSOs).

**Figure 3.**
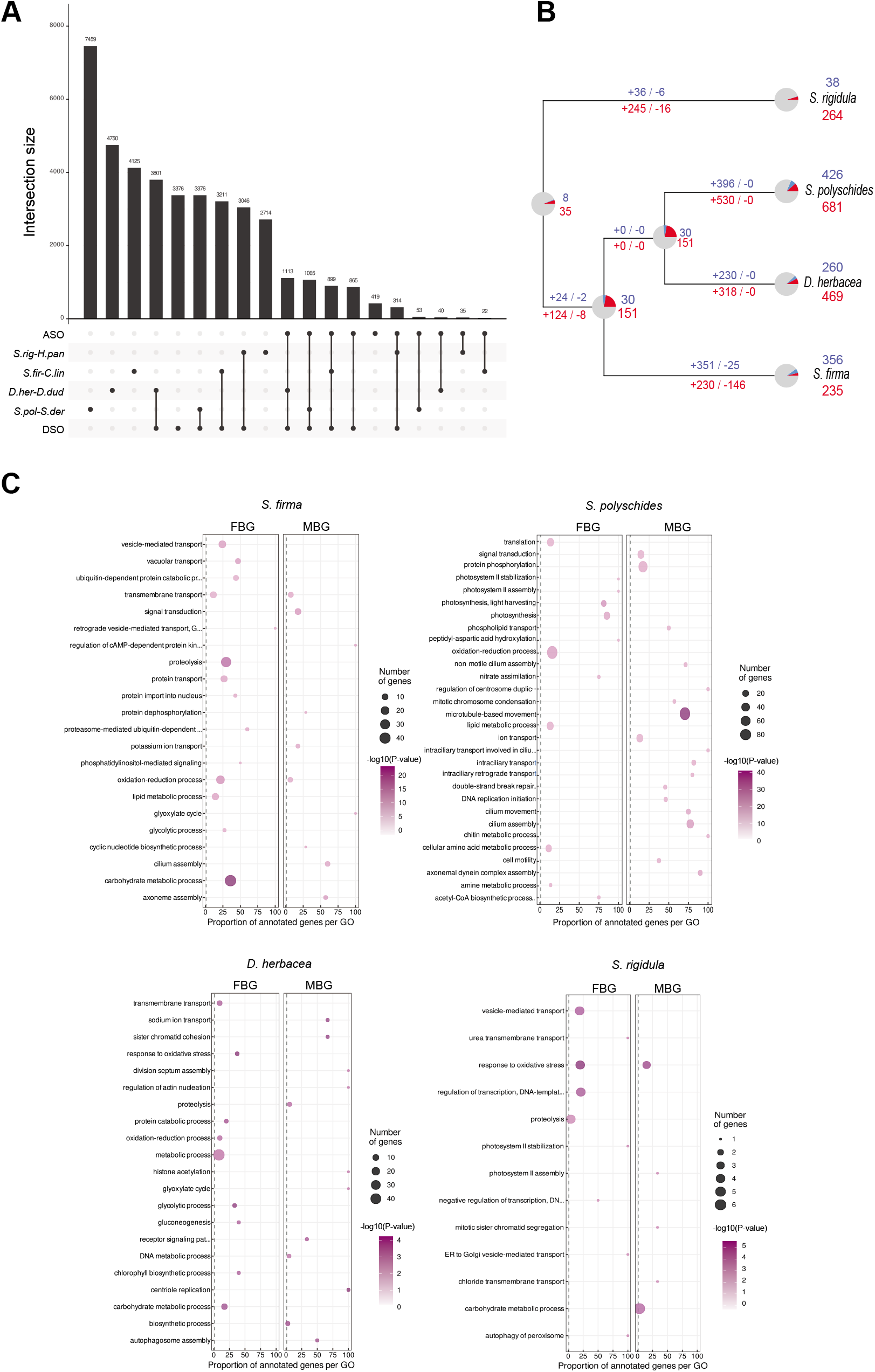
Single copy orthologs gene sets and sex-biased genes ancestry. A) Representation of intersects across single-copy ortholog gene sets (i.e., the four PSO, DSO and ASO) using UpSetR. Bars represent the number of genes in the intersect represented below the histogram. PSO: pairwise single-copy orthologs. DSO: dioicous single-copy orthologs. ASO: All species single-copy orthologs. B) Reconstruction of ancestral sex-biased gene sets across the four dioicous species. The number of inferred sex-biased genes (female-bias in red, male-bias in blue) at ancestral nodes as well as the inferred gain and loss of sex-biased genes along branches are displayed. C) Enriched GO-terms associated with sex-biased genes from each

We then used maximum likelihood approaches to infer the ancestral states of sex-biased gene expression across these dioicous species (Figure 3B). Our analysis identified very few genes that were predicted to be ancestrally sex-biased, with the vast majority having evolved sex-bias at some point along the branches. Among the 2,116 sex-biased DSOs in at least one species, only 43 (2.03%) were inferred to be sex-biased in the last common ancestor of the four brown algal species (Figure 3B). Accordingly, no DSOs were consistently sex-biased across the four species (not different from what is expected by chance, exact test multi-set intersection *P* = 0.506). A total of 139 OGs exhibited a bias in one species that was inconsistent with the direction of bias observed in at least one other species (Table S4).

Although the above analysis showed that sex-bias genes were not conserved among the four species, we examined if sex-biased genes in different species were involved in similar functions, by comparing gene ontology (GO) terms of sex-biased genes across species using Blast2Go ^28^. We detected significant enrichment of GO terms for biological processes related to ‘microtubule’, ‘ion transport’ and ‘cilium’ consistently for male-biased genes across all dioicous species. Conversely, the sets of female-biased genes of all four species were enriched for GO terms such as ‘photosynthesis’, metabolism, ‘oxidation/reduction processes’ (Figure 3C, Table S5).

Taken together, our results indicate that whilst the different species do not share the same sex-biased gene set, males and females across these brown algae display a striking convergence in terms of sex-biased gene functions.

### Sex-biased gene expression fate during transition to monoicy

To study changes in sex-biased gene expression that accompany the transition from dioicy to monoicy, we first identified single-copy orthologous genes for each of the four dioicous-monoicous sister species pairs (pairwise single-copy OGs; PSOs, Figure 2A). We were able to infer between 6,109 and 11,953 PSOs for each of the four pairs of species (Figure 2A; Tables S6-S10). PSOs were classified as being sex-biased or unbiased by comparing male and female expression in each dioicous species (FDR < 0.05, FC> 2). We then examined the patterns of expression of male-, female- and unbiased PSOs in dioicous males and females and in the corresponding monoicous species.

In three out of the four species pairs, the levels of expression of sex-biased genes in the monoicous species were similar to the values measured for orthologues in females of the corresponding dioicous species (Figure 4A). In these three pairs, male-biased genes were downregulated in the monoicous species compared with males, and they displayed similar expression levels to male-biased genes in females of the dioicous species, suggesting that de-masculinisation of gene expression of the monoicous species counterpart occurred frequently. Female-biased genes were expressed at similar levels in *S. firma* and *D. herbacea* females compared with the corresponding monoicous species. In the *S. polyschides-S. dermatodea* pair of species, female-biased genes had a similar pattern in males and monoicous. In the *S. rigidula/H. paniculata* species pair, no significant difference was detected between the expression of sex-biased and unbiased genes between the two species. Note, however, that results for *S. rigidula/H. paniculata* were more difficult to interpret, as the low number of sex-biased genes precluded robust statistical analysis.

**Figure 4.**
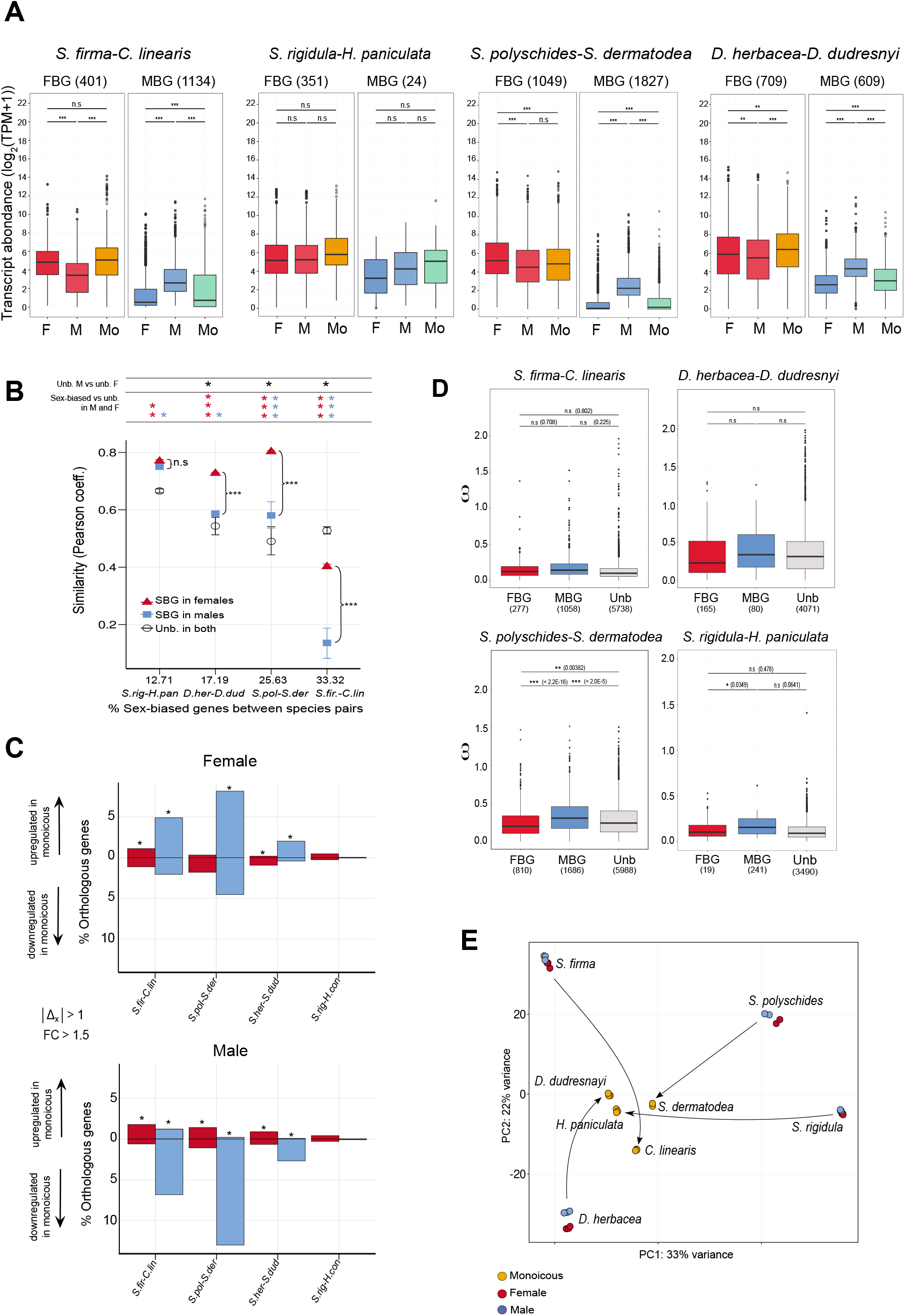
Evolution of sex-biased genes during transitions to monoicy. A) Comparison of gene expression levels within species pairs, in log2(TPM+1), using PSO gene sets. F: females. M: males. Mo: monoicous. The number of female-biased (FBG) and male-biased genes (MBG) among PSO are displayed. Statistical tests are permutation t-tests using 100,000 permutations. B) Comparisons of similarity index values (Pearson coefficients) between expression profiles (in log2(TPM+1)) of genes of dioicous and monoicous species within species pairs (PSO). Similarity index are represented separately for sex-biased genes in females (red) and in males (blue), as well as for unbiased genes averaged across sexes (black). Pearson coefficients were plotted for each species pair in increasing order of the proportion of SBG among expressed genes of dioicous species. Stars in the top pannel represent significant differences between Pearson coefficients, taking into account the correlations between compared gene sets, using the cocor package in R. Red and blue stars indicate a significant difference between females (red stars) or males SBG (blue stars) coefficients with unbiased genes coefficients. Top panel black stars indicate a significant difference of Pearson coefficients of unbiased genes between males and females. Significant differences of coefficients between sex-biased genes in females and males are indicated directly on the plot. * : 0.01 < P < 0.05; ** : 0.001 < P < 0.01; *** : P < 0.001. C) Fraction of female-biased genes (red) and male-biased genes (blue) with an absolute value of Δ_X_ >1 and a fold-change > 1.5, calculated within species pairs (on PSO). Downregulated genes in the monoicous species are represented below the y=0 line (Δ_X_ < −1), upregulated genes in the monoicous species are represented above the y=0 line (Δ_X_ >1 ). Stars indicate a significant over-representation of female-biased (red star) or male-biased genes (blue star) with an absolute Δ_X_ >1 compared with the proportion of unbiased genes with Δ_X_ >1, tested using Fisher exact tests. A reminder of differences between male and female gamete size within species pair is sketched below the x-axis. D) Sequence divergence, measured as d_N_/d_S_ (ω), between dioicous and monoicous species calculated within species pairs (PSO). Statistical tests are permutation t-tests using 100,000 permutations, p-values are displayed in parentheses. * 0.01 < P < 0.05; ** 0.001 < P < 0.01; *** P < 0.001. E) Principal component analysis (PCA) plot of all the RNA-seq samples, using ASOs. Monoicous species are plotted in brown, female samples in red and male samples in blue.

We next investigated the gene expression profiles of monoicous species in order to test whether their transcriptional patterns resemble those of their male or female dioicous counterparts. We computed the Pearson product-moment coefficient of regressions of gene expression profiles (in log2(TPM+1)) of males or females compared with that of the monoicous species within each species pair. We compared Pearson correlation coefficients for both sex-biased genes and unbiased genes in both males or females, considering sex-biased genes in males and females as independent groups. We also compared the correlations of expression profiles with the orthologs of sex-biased and unbiased genes in the monoicous species, separately for males and females. These groups of sex-biased versus unbiased genes being expressed within the same individuals, we considered them as dependant groups in the *cocor* package (Diedenhofen & Musch, 2015). Altogether, these analyses indicated that, with the exception of the *S. rigidula-H. paniculata* species pair, the gene expression profiles of the monoicous species were significantly more similar to those of females than they were to male profiles (Figure 4B). Moreover, the close association between female and monoicous expression profiles was observed for both sex-biased and unbiased genes (Figure 4B, black asterisks at the top).

Interestingly, with the exception of the Ectocarpales species pair (*S. firma*-*C. linearis*), sex-biased gene expression profiles diverged significantly less from the monoicous species than did that of the unbiased genes. We also noted that the highest similarity indexes for within species pairs were found for the species with the lowest level of sex-biased gene expression (*S. rigidula*), and the lowest similarity was observed for *S. firma*, the species with the highest level of sex-biased gene expression (Figure 4B).

Taken together, the above observations suggest that gene expression profiles of monoicous species tend to be more closely related to the females of the related dioicous species, and this similarity appears to be driven by sex-biased genes, in particular male-biased genes. The tendency to reproduce the female transcriptome in the monoicous species was repeatable in independent transitions to co-sexuality.

### Is selection involved in gene expression changes during transition to monoicy?

To examine whether changes in gene expression during transition to co-sexuality were the result of selective or neutral forces, we computed the degree of directional selection using Δx. This parameter evaluates the divergence in expression level in relation to the variation in expression level seen across replicates ^5,26,29^. We computed Δx of the PSO sets, separately for each pair of species and reported the proportions of orthologs with an absolute Δx > 1, i.e., orthologs whose expression shift is attributable to directional selection.

Depending on the species pair, between 10.8% and 50% of unbiased genes exhibited expression shifts attributable to selection (|Δ_x_|>1) (Table S11). We then asked whether male- and female-biased genes were preferentially concerned by adaptive expression shifts during transitions to monoicy compared with unbiased genes. Fisher’s exact tests showed that for three out of four species pairs, male-biased genes were indeed more likely to display |Δ_x_|>1 than unbiased genes. This was also the case for female-biased genes in *S. polyschides-S. dermatodea* pair of species (Fisher exact tests, *P*<2.2 ×10^−16^ in both sexes) and *D. herbacea-D. dudresnayi* pair (Fisher exact tests, *P* = 3.9 x10^−3^ and *P* = 1.8 ×10^−5^ in females and males, respectively). In *S. polyschides-S. dermatodea* and *S. rigidula-H. paniculata*, female-biased genes showed lower levels of adaptive evolution of expression compared with unbiased genes (Table S11, Figure 4C). Taken together, our observations indicate that male-biased genes preferentially exhibit a shift in expression during the transition to monoicy that may be explained by directional selection.

We also assessed if evolution of gene expression during the transition to monoicy has been driven by DNA sequence evolution, by using measures of sequence divergence (*d*_N_/*d*_S_). We computed *d*_N_/*d*_S_ for male-biased, female-biased and unbiased genes for each of the dioicy-monoicy species pairs. For all four pairs, male-biased genes consistently exhibited higher evolutionary rates than female-biased and unbiased genes, although this difference was significant only for the *S. polyschides-S. dermatodea* species pair (Figure 4D, Table S12). As this is the ‘youngest’ species pair (Figure 1), it appears that the level of sequence divergence during transition to monoicy is not associated with the age of transition.

Taken together, our observations indicate that shifts from dioicy to monoicy involved modifications to transcriptional patterns (expression divergence) mostly at male-biased genes that were likely driven by selection but also coding sequence evolution.

### Convergent gene expression changes associated with transition to monoicy

In order to assess the extent to which gene expression changes occurring during the transition to monoicy were shared across the four species pairs, we focused on the single-copy orthologs across the eight species, herein termed ‘All Single-copy Orthologs’ (ASO). We found a total of 1,708 ASO (following the same methods as for DSO, see methods).

Among the 1,708 ASOs, 718 were sex-biased in at least one dioicous species (Tables S13, S14). Sex-biased genes were not over-represented among ASOs (Fisher exact test, *P* = 0.097). Sixty one percent of the ASOs (1,043 out of 1,708) exhibited a conserved pattern of expression across all monoicous species compared to the dioicous species. This proportion was significantly different from what was expected by chance (permutation tests, *P* = 0.0255, 10,000 permutations) suggesting convergent gene expression changes during transition to monoicy across all studied pairs of species. Decomposition of variance components for the 1,708 ASOs detected a clear pattern of grouping of monoicous species, further illustrating the extensive convergence of gene expression during the transition from dioicy towards monoicy (Figure 4E). Functional analysis of genes that are convergently expressed during the transition to monoicy highlighted terms such as nucleic acid metabolic processes and transmembrane transport (Figure S2).

About half (527) of the 1,043 genes that were consistently differentially expressed in monoicous versus dioicous species had a |*Δ*_*X*_| >1, which is significantly more in proportion than among the rest of the ASO (290 genes with |*Δ*_*X*_| >1 among 665 ASO, Fisher exact test *P* = 0.00543). This observation indicates that convergent gene expression changes may be associated with directional selection during the switch to monoicy.

We next tested whether sexual selection potentially occurring in males and females of dioicous species would be relaxed in monoicous individuals. This would translate by a reduction of purifying selection resulting in increased sequence divergence (increased *d*_*N*_/*d*_*S*_). Convergent genes (i.e., genes exhibiting a convergent pattern of gene expression in monoicous species) tended to exhibit faster divergence rates compared with non-convergent genes although the difference was not significant (permutation *t*-test, *P* = 0.0566). Noteworthy, male-biased (but not female-biased) genes showed significantly higher *d*_N_/*d*_S_ than unbiased genes (Table S15).

A likelihood ratio test of branch models (after Benjamin-Hochberg correction for multiple testing), identified 689 orthologs under positive selection on monoicous branches, 404 of which exhibited convergent gene expression changes. Orthologs under positive selection were over-represented among genes with convergent gene expression (Fisher exact test, *P* = 0.025). Taken together, these observations suggest that directional selection plays a role in driving changes in expression patterns during transitions to co-sexuality.

## Discussion

### Phenotypic sexual dimorphism and sex-biased gene expression are uncoupled in brown algae

Although morphological and physiological differences between males and females are ultimately due to divergences between sex chromosomes in species with genetic sex determination (Bachtrog et al. 2014), the majority of morphological sexual dimorphism is thought to be associated with autosomal sex-biased gene expression ^3–5^. Thus, it would be expected that species showing more prominent differences in morphology between male and female would also be characterised by high levels of sex-biased gene expression, as has been shown to be the case in birds ^27^. Our study, in contrast, revealed no correlation between the level of sex-biased gene expression and the degree of phenotypic sexual dimorphism in the brown algae studied here. Therefore, the link between gene expression evolution and sexual selection is uncertain for these organisms, and may reflect a lower degree of sexual selection in the brown algae compared with animals. Brown algae have relatively low levels of sexual dimorphism ^9,19^ and are broadcast spawners so the opportunities for mate choice and/or mating competition are mainly constrained to interactions involving male and female gametes and not gametophytes ^30^. Consistent with the idea that gamete sexual selection may occur (and perhaps not so much gametophyte sexual selection), it has been shown recently that in the absence of males, female gametes of brown alga populations lose their sexual morphological characteristics, e.g. female gametes produce lower levels of pheromone and engage in parthenogenesis more rapidly ^31^.

### Sex-biased genes have high turnover but exhibit functional convergence

Although dioecy is predicted to be the ancestral sexual system in brown algae ^22^ our results clearly indicate that sex-bias in the expression of individual genes is neither ancestral nor convergent. We found a very limited level of shared (ancestral) sex-biased gene expression across the studied brown algal species, and instead our data is consistent with lineage-specific recruitment of sex-biased genes. Our observations emphasize therefore a substantial turnover of sex-biased expression among brown algal genes.

Although the dioicous brown algal species studied here shared very few sex-biased genes, we found a remarkable level of convergence in terms of sex-biased gene function. These include biological functions that were previously found to be enriched in *Ectocarpus* gametophytes ^9,32^ further underscoring the conservation of sex-biased gene function and supporting primary sexual dimorphic roles. Considering that brown algae share an ancestral sex chromosome, and that genes within the non-recombining sex determining region play a role in sex ^33^, one possibility is that sexual characteristics in these UV systems mainly involve genes within the SDR together with a relatively limited number of autosomal genes involved in primary sexual dimorphisms. In other words, differences between sexes arise mainly from the different physiological processes directly linked to the production of male or female gametes rather than extensive sexual selection, sexual specialization and/or sexual antagonism (i. e, secondary sexual dimorphism) ^6^.

### Fate of sex-biased gene expression during the transition to monoicy

Our sampling of species distributed across the brown algae phylogeny, associating pairs of related dioicous and monoicous species, allowed us to trace the fate of sex-biased gene expression during independent events of transition from dioicy to monoicy. With the exception of one species pair, sex-biased genes exhibited adaptive expression shifts during the transition to monoicy. Male-biased genes, specifically, were the main drivers of gene expression changes during the transition to monoicy, while unbiased genes exhibited limited changes in pattern of expression with the switch in sexual system. In the model brown alga *Ectocarpus*, RNA-seq analysis of multiple tissues and life cycle stages indicated that sex-biased genes have restricted patterns of expression, which is a proxy for limited pleiotropy ^9^. Pleiotropy is known to restrict gene evolution, imposing stricter functional constraints on pleiotropic genes^34,35^. The reduced pleiotropy of sex-biased compared with unbiased genes may increase their propension to adaptively shift towards their evolved optimal expression profile during evolutionary transitions, in this case the transition to monoicy ^4,35,36^.

Sex-biased genes in dioicous brown algae such as *Ectocarpus* sp. typically display high evolutionary rates compared to unbiased genes due either to directional selection or relaxed purifying selection ^9^. With the transition to monoicy, increased relaxation of sex-specific purifying selection acting on sex-biased genes may be expected, leading to increased rates of sequence evolution. Accordingly, male-biased genes for all species pairs presented faster evolutionary rates (although not significant for all species) during the switch to monoicy, compared with female-biased or unbiased genes. This observation points to a shared process of sexually antagonistic selection within dioicous species, especially in males, that allowed for faster evolutionary rates of male-biased genes when relaxed during the transition from dioicy to monoicy.

### Convergent changes associated with the breakdown of dioicy and origin of monoicy

Convergent evolution, where a similar trait evolves in different lineages, provides an opportunity to study the repeatability of evolution. In the brown algae, co-sexuality has repeatedly emerged from uni-sexual ancestors ^22^. Strikingly, we found that more than half (61%) of the orthologs across the four pairs of species displayed similar expression shifts concomitant with the transition to monoicy, indicating that common, independently acquired mechanisms are associated with co-sexuality. Remarkably, a substantial number of these convergent genes (38.7%) were under positive selection, underlying the idea that convergent changes associated with the shift of sexual system may be driven by comparable evolutionary pressures across these distant species.

In our study, the expression profiles of gametophytes of all four monoicous species resembled those of the female gametophytes of their dioicous counterpart. Moreover, sex-biased genes tended to maintain the level of expression they had in dioicous species, suggesting that they retained their ancestral function in the context of derived monoicy. When their expression shifted, sex-biased genes, and especially male-biased genes showed signs of selection acting on their expression level to a greater extent than it acted on unbiased genes. Together, our results demonstrate that common mechanisms underlie the transition to monoicy across distant brown algal lineages and suggest that independent events of loss of dioicy may have involved acquisition of genes related to male development by a female gametophyte. The work presented here establishes therefore a framework for understanding at the genomic level how co-sexual systems arise from ancestral haploid UV sexual systems.

## Materials and methods

### Sample preparation, RNA extractions and sequencing

The algal strains used and sequencing statistics and BioProject accession number are listed in Table S1. Gametophytes of all eight species were cultured at 13 °C in autoclaved natural sea water (NSW) supplemented with half-strength Provasoli solution (PES; ^37^) with a light:dark cycle of 12:12 h (20 μmol photons m^−2^ s^−1^) using daylight-type fluorescent tubes ^38^. All manipulations were performed under a laminar flow hood in sterile conditions. Immature gametophytes (i.e., absence of sex-specific reproductive structures, oogonia or antheridia) of each strain were frozen in liquid nitrogen and kept at −80C until RNA extraction.

RNA from male and female pools was extracted from triplicate samples (each containing at least 800 individual gametophytes; except for *S. polyschides* and *S. dermatodea* where two replicates were used) using Qiagen RNA extraction Plant Mini kit. RNA quality and quantity were assessed using an Agilent 2100 bioanalyzer, associated with an RNA 6000 Nano kit. For each replicate, the RNA was quantified and cDNA was synthesized using an oligo-dT primer. The cDNA was fragmented, cloned, and sequenced by Fasteris (CH-1228 Plan-les-Ouates, Switzerland), using Illumina Hi-seq2000 for Saccorhiza and Desmarestia species; by Genome Quebec using an or Nextgen6000 for Halopteris and Chordaria species; and by Genoscope using Illumina Hi-seq 4000 for Sphacelaria and Sphaerotrichia species (see Table S1 for details).

### Transcriptome assemblies and gene set predictions

Predicted gene sets were constructed for each species base on genome and transcriptome assemblies. In order to filter out potential contamination, first round assembled contigs were blasted against the NCBI non-redundant (nr) protein database using diamond v 0.9.21 ^39^ and reads that mapped on contigs with non-eukaryotic taxons were removed using blobtools v 1.0.1 ^40^. *De novo* transcriptomes were assembled using Trinity (*Saccorhiza polyschides*, *Saccorhiza dermatodea*, *Desmarestia dudresnayi, Desmarestia herbacea* female, *Halopteris paniculata*, *Sphacelaria rigidula*) or rnaSPADES v 3.12.0 (*Chordaria linearis*, *Sphaerotrichia firma*) with kmer size of 55.

All genomes were soft-masked using Repeatmasker v 4.0.9 after building a *de novo* transposable elements and repeats database with RepeatModeler v 1.0.8 ^41^. BRAKER2 ^42^ and PASA (for *Desmarestia herbacea* ^43^, using input predicted protein from the reference species *Ectocarpus* sp. (EctsiV2_prot_LATEST.tfa ^44^, were used to predict gene sets used for all downstream analyses.

The final assemblies are available in NCBI (BioProject accession number PRJNA733856). Transcriptome completeness was assessed using BUSCO v3 eukaryote gene set as reference (Odb9). Transcripts that had DNA data support for only one sex (potentially sex-linked) were tested with PCR using at least 4 males and 4 females per species and removed from the sex-biased gene analysis. PCR primers are detailed in Table S16.

### Expression quantification and identification of sex-biased genes in dioicous species

RNAseq reads adaptors were trimmed using trimmomatic v0.38 ^45^ which was also used for read-quality filtering: reads were removed if the leading or trailing base had a Phred score <3, or if the sliding window Phred score, averaged over four bases, was <15. Reads shorter than 36 bases were discarded (as well as pair of reads, if one of the pair was <36 bases long). Trimmomatic-processed RNAseq reads from each library were used to quantify gene expression with kallisto v 0.44.0 ^46^ using 31 bp-long kmers and predicted transcript of each species. RNAseq libraries were composed of stranded (--fr-stranded or --rf-stranded option) single-end reads (--single option) or paired-end reads (Table S1). A gene was considered expressed in a given species and/or a given sex when at least one library displayed an expression level (in TPM) above the 5^th^ percentile of TPM distribution across all genes and libraries within a species and sex. Following 47transcript abundances were then summed up within genes and multiplied by the total library size, using the tximport package^25^ to obtain the expression level for each gene in transcripts per million reads (TPM).

Estimates of sex-biased gene expression in dioicous species were obtained using read count matrices as input for the DESeq2 package (Love et al., 2014) in R 3.6.3. *P-*values were corrected for multiple testing using Benjamini and Hochberg’s algorithm in DESeq2, applying an adjusted *P*-value cut-off of 0.05 for differential expression analysis. Genes with a minimum of 2-fold change expression level between sexes were retained as sex-biased.

### Quantification of phenotypic sexual dimorphism

Individual gametophytes from each of the strains were isolated in sea water and observed using an inverted transmitted light microscope DMi8 (Leica) with the LAS X software. Between 269 and 556 cells (348 cells on average per sex and per species) across five different gametophytes per species were individually measured using Fidji ^48^. We used *t*-tests to compare cell length between groups. The difference in mean cell length between sexes of dioicous species was computed and used as a proxy for phenotypic sexual dimorphism. To investigate the relationship between phenotypic sexual dimorphism and extent in sex-biased expression, phenotypic dimorphism was regressed against the fraction of sex-biased genes in R.

### Orthology and evolutionary rates within species pairs

We inferred pairwise single-copy orthologs (PSO) within the four species pair using Orthofinder with default parameters ^49^. We used kallisto v 0.44.0 to quantify expression levels for PSO within species pairs. In order to infer the potential role of selection in expression changes between dioicous and monoicous species we computed *Δ*_*X*_. To summarize we calculated *Δ*_*X*_ = *d / r* with *d* and r respectively given by:

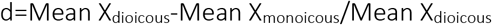

and

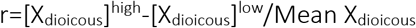

where X is the expression level measured in TPM, ‘High’ and ‘Low’ represent the maximum and minimum values. Fisher exact tests were computed to detect whether female-biased genes (FBG) and male-biased genes (MBG) were more likely to show an absolute value of *Δ*_*X*_ > 1 compared to unbiased genes.

Orthologous proteins between species pairs were aligned with MAFFT v7.453 ^50^ the alignments were curated with Gblocks v0.91b ^51^ and back-translated to nucleotides using translatorX ^52^. We used these nucleotide alignments as input for phylogenetic analysis by maximum likelihood (PAML4, CodeML, ^53^) to infer pairwise *d*_N_/*d*_S_ (ω) with F3×4 model of codon frequencies. We retained orthologs with 0 < *d*_*S*_ < 2 as valid for further analysis. We compared species and sexes evolutionary rates separately for female-biased, male-biased and unbiased genes, using permutation *t*-tests in R with 100,000 permutations.

### Evolution of sex-biased gene expression

We inferred a single orthologous gene set for the four dioicous species (DSO) using Orthofinder with default parameters. Following the methods used in^54^ we included in the DSOs the orthogroups genes that were 1:1:1:0, likely due to situations in which a single-copy ancestral gene was lost in a single species. To further account for gene prediction errors, we also included orthogroups with a single species presenting two-genes that aligned on more than 60% of their length as duplicate genes.

A well resolved phylogeny of the Pheaophyceae was used as reference gene tree ^24^ to infer where sex-biased gene expression evolved along the phylogenetic tree. We coded DSO as either male-biased, female-biased or unbiased for each species and used the ape package ^55^ in R to reconstruct the discrete ancestral state *via* maximum likelihood. Proportions of ancestral genes in each category were plotted as pie-charts on tree nodes and gain/loss of bias were reported on each branch. We further tested the significance of overlap between sex-biased genes identified within dioicous species with exact multi-set intersection test implemented in the SuperExactTest package v 1.0.7 in R ^56^.

We inferred expression profile similarity index between monoicous species and males and females of dioicous species within pairs as the Pearson correlation coefficient of PSO expression levels in log2(TPM + 1). This analysis was performed for all expressed genes, and separately for MB, FB and unbiased genes. We compared Pearson coefficients of regression within each species pair, using the cocor package ^57^, considering gene expression profiles of males and females as independent gene sets. We also compared SBG with unbiased genes within sexes, considering these gene sets as dependent. We report the *P*-value based on Fisher’s *z* or, when possible, Silver, Hittner and May’s modification of Dunn and Clark’s *z.* Pearson’s coefficients were plotted for each species pair.

### Convergent expression changes

Convergent changes associated with transitions to monoicy were investigated on single-copy orthologs inferred across the eight studied species (termed ‘All Single-copy Orthologs’, ASO) following the same methods as those used for the DSOs. Using this data set, we quantified gene expression with kallisto as described above, and DESeq2 was used to infer orthologs significantly affected by sexual system but not species pair (lfcShrink with “ashr” method, sexual system contrast; Stephens, 2016). Significance of the number of convergent expression changes was tested with permutation tests (100,000 permutations). We used the ComplexHeatmap package ^58^ in R to visualize gene expression for each replicate. Orthogroups with inconsistent sex-bias across different species (n=139) were removed from the dN/dS analysis of convergent gene evolution.

Intersects between genes across PSO, DSO and ASO were represented using the UpSetR package v1.4.0 ^59^.

### ASO Evolutionary rates

Following the same process described for pairwise orthologs, we aligned and studied molecular sequence divergence for all species orthologs (ASO) using CodeML. We used a ‘two-ratio’ branch model (model = 2, Nssites = 0) to specifically study divergence on monoicous branches (foreground branches). We compared ω values separately between sex-biased (male-biased and female-biased genes) and unbiased genes with permutation *t*-tests (10,000 permutations). We also ran two branch-site models in PAML to detect positive selection in foreground branches (model=2, Nssite=2, ω=1 fixed (Null) or allowed to vary). Likelihood-ratio tests were used to compared the model of selection with the null model in order to detect orthologs with sites under positive selection in the monoicous branches. LRT *P*-values were corrected for multiple testing using Benjamini and Hochberg’s algorithm^60^.

### Functional annotation analysis

Predicted genes and orthogroups were blasted against the NCBI non-redundant (nr) protein database with blast (v2.9.0). Functional annotation was performed using BLAST2GO ^28^, as well as the InterProScan prediction of putative conserved protein domains ^61^. Gene set enrichment analysis was carried out separately for each geneset using Fisher’s exact test implemented in the TopGO package, with the *weight01* algorithm ^62^. We investigated enrichment in terms of biological process ontology and reported significant GO-terms with *P*-value < 0.01.

All statistical analyses were performed in R 3.6.3, plots were produced with ggplot2 in R (https://ggplot2.tidyverse.org/).

## Acknowledgments

We would like to thank J.R. Pannell for valuable comments on the present study, I. Theodorou for help with microscopy, A. F. Peters for help with algal cultures and T. Broquet for comments and advice on statistical methods. This work was supported by an ERC grant to SMC (grant agreement 864038) and the France Génomique National infrastructure project Phaeoexplorer (ANR-10-INBS-09).

## Data

Raw reads have been deposited in the SRA. Accession codes are given in Table S17.

## Supplemental Figures

**Figure S1.**
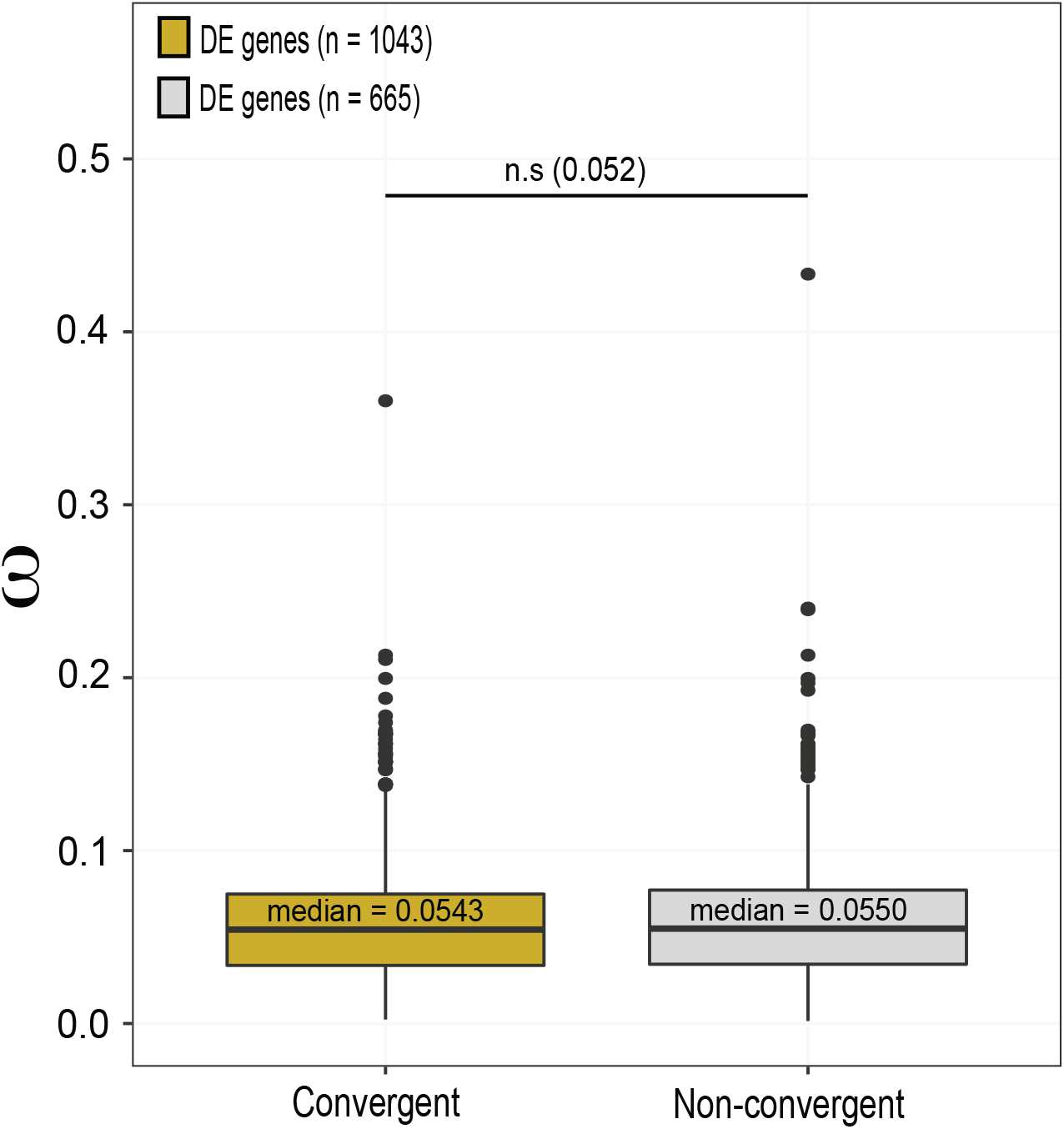
Sequence divergence, measured as d_N_/d_S_ (ω), between convergent and non-convergent genes across all species pairs (ASO).

**Figure S2.**
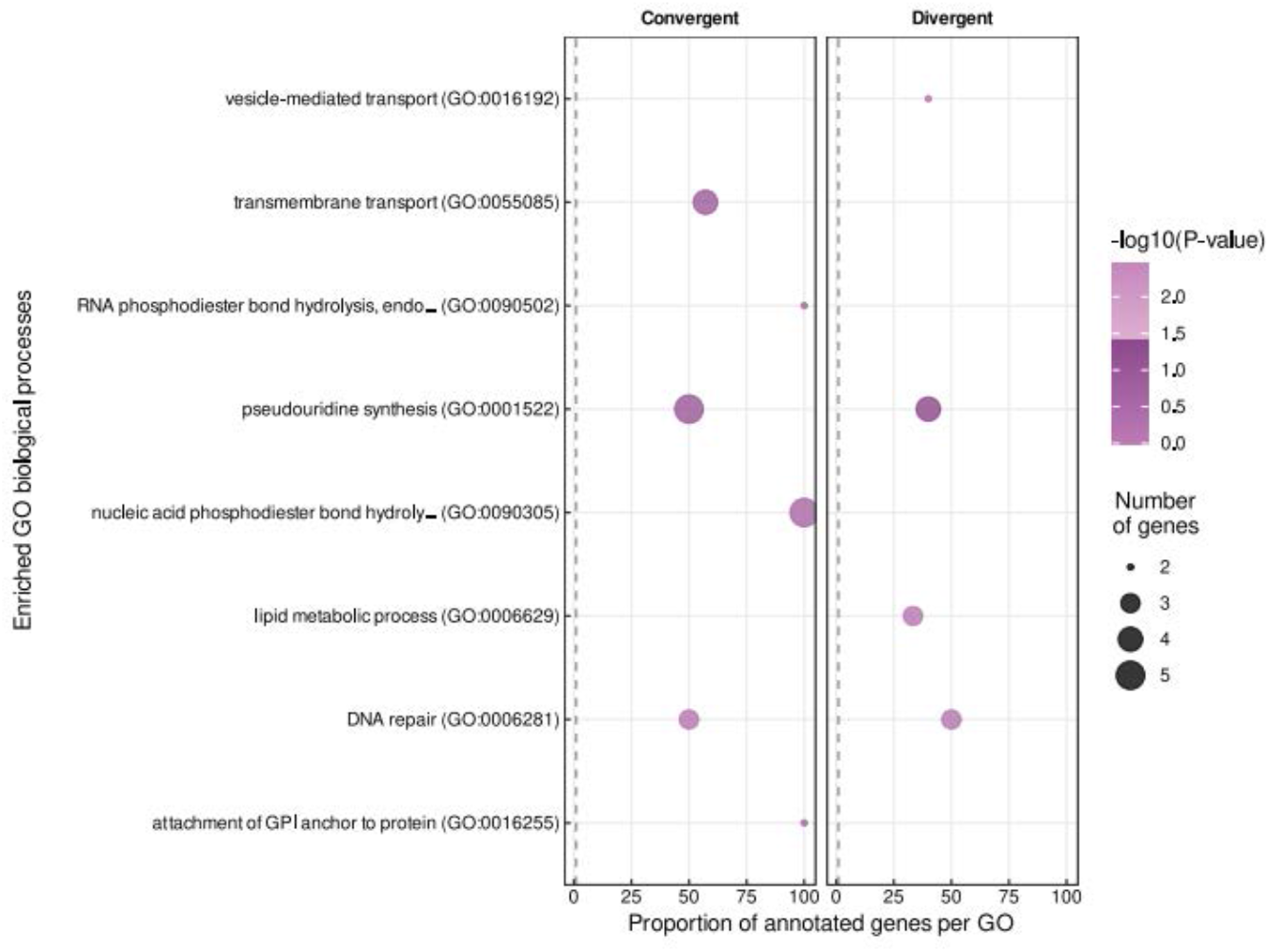
Functional analysis (enriched GO terms for biological processes) of genes that are convergently or divergently expressed in dioicous versus monoicous species

## Table Legends

Table S1. Brown algae species used in the study and summary of gene expression data sets.

Table S2. Summary statistics of sex-biased gene expression. Fractions of sex-biased genes are calculated over the number of expressed genes in the sex of bias.

Table S3. Summary of sexually dimorphic traits in the four dioicous brown algae species investigated in the present study.

Table S4. Orthogroups belonging to the dioicous single-copy orthologs geneset (DSO) and their corresponding gene and bias per species.

Table S5. Description of GO-term enrichment of sex-biased genes identified in each dioicous species. *P*-values correspond to Fisher exact tests (FDR < 0.05).

Table S6. Summary description of all pairwise single-copy orthologs (PSO).

Table S7. Orthogroups belonging to the pairwise single-copy orthologs genes et (PSO) within the Tilopteridales species pair. Mean expression level in TPM across replicates as well as the bias status in the dioicous species are reported.

Table S8. Orthogroups belonging to the pairwise single-copy orthologs gene set (PSO) within the Desmarestiales species pair. Mean expression level in TPM across replicates as well as the bias status in the dioicous species are reported.

Table S9. Orthogroups belonging to the pairwise single-copy orthologs geneset (PSO) within the Sphacelariales species pair. Mean expression level in TPM across replicates as well as the bias status in the dioicous species are reported.

Table S10. Orthogroups belonging to the pairwise single-copy orthologs geneset (PSO) within the Ectocarpales species pair. Mean expression level in TPM across replicates as well as the bias status in the dioicous species are reported.

Table S11. Summary statistics of Δ_X_ within the four species pairs. Male-biased genes (MBG) and female-biased genes (FBG) that were significantly more likely or less likely to present |Δ_X_| > 1 were highlighted in green and purple, respectively (Fisher’s exact tests).

Table S12. *P*-values of permutation *t*-tests (100,000 permutations) of sequence divergence data (dN/dS) calculated within species pair, between female-, male-biased and unbiased genes.

Table S13. Summary description of all single-copy orthologs (ASO)

Table S14. Orthogroups belonging to the all single-copy orthologs geneset (ASO). Mean expression level in TPM across replicates as well as the bias status in the dioicous species are reported.

Table S15. *P-*values of permutation *t*-tests (10,000 permutations) of sequence divergence data (dN/dS), calculated specifically for monoicous branches (branch model) across ASOs, between female-, male-biased and unbiased genes. Significant difference of divergence with unbiased genes are put in bold.

Table S16. Primers used to test candidate sex-linked contigs in the different brown algal species.

Table S17. Accession references.

